# Chromosome-level assembly of *Drosophila bifasciata* reveals important karyotypic transition of the X chromosome

**DOI:** 10.1101/847558

**Authors:** Ryan Bracewell, Anita Tran, Kamalakar Chatla, Doris Bachtrog

**Affiliations:** Department of Integrative Biology, University of California Berkeley, Berkeley, CA 94720, USA

## Abstract

The *Drosophila obscura* species group is one of the most studied clades of Drosophila and harbors multiple distinct karyotypes. Here we present a *de novo* genome assembly and annotation of *D. bifasciata*, a species which represents an important subgroup for which no high-quality chromosome-level genome assembly currently exists. We combined long-read sequencing (Nanopore) and Hi-C scaffolding to achieve a highly contiguous genome assembly approximately 193Mb in size, with repetitive elements constituting 30.1% of the total length. *Drosophila bifasciata* harbors four large metacentric chromosomes and the small dot, and our assembly contains each chromosome in a single scaffold, including the highly repetitive pericentromere, which were largely composed of Jockey and Gypsy transposable elements. We annotated a total of 12,821 protein-coding genes and comparisons of synteny with *D. athabasca* orthologs show that the large metacentric pericentromeric regions of multiple chromosomes are conserved between these species. Importantly, Muller A (X chromosome) was found to be metacentric in *D. bifasciata* and the pericentromeric region appears homologous to the pericentromeric region of the fused Muller A-AD (XL and XR) of *pseudoobscura*/*affinis* subgroup species. Our finding suggests a metacentric ancestral X fused to a telocentric Muller D and created the large neo-X (Muller A-AD) chromosome ∼15 MYA. We also confirm the fusion of Muller C and D in *D. bifasciata* and show that it likely involved a centromere-centromere fusion.

## INTRODUCTION

Recent advances in DNA sequencing technology have dramatically improved the quality and quantity of genome assemblies in both model and non-model species. Long-read sequencing technologies (e.g., PacBio and Nanopore) combined with long-range scaffolding information generated through chromatin conformation capture methods such as Hi-C (Lieberman-Aiden et al. 2009) or Chicago (Putnam et al. 2016) can produce assemblies of unprecedented length and accuracy. However, there are still relatively few assemblies that traverse through megabase-long stretches of highly repetitive sequence, thereby limiting our understanding of the evolution of pericentromere/heterochromatic regions of the genome and the genes, satellites, and transposable elements that inhabit them (Chang et al. 2019, Miga 2019).

Drosophila has been at the forefront of genetics and genomics research for over a century and new chromosome-level assemblies are now becoming available for several non-model species (Mahajan et al. 2018, Miller et al. 2018, Bracewell et al. 2019, Karageorgiou et al. 2019, Mai et al. 2019). Recent comparative genomic analysis in the *Drosophila obscura* group has revealed extensive karyotype evolution and turnover of centromeric satellites that alters chromosome morphology (Bracewell et al. 2019) (Figure 1). Unfortunately, our understanding of karyotype and genome evolution is currently limited because no high-quality assembly of a species from the *obscura* subgroup is available (Figure 1). Given the phylogenetic placement of *D. bifasciata* (Figure 1) and its putative chromosomal configuration (Buzzati-Traverso and Scossiroli 1955, Moriwaki and Kitagawa 1955), it is an important species for reconstructing karyotype evolution in the *obscura* group for several reasons. First, a high-quality *D. bifasciata* genome assembly allows us to better understand the emergence of metacentric chromosomes and determine if metacentric pericentromeres are conserved over evolutionary time (Figure 1). Second, the configuration of the Muller A chromosome (the ancestral X chromosome in Drosophila) is particularly interesting, since it became fused to Muller D in some members of the *obscura* group ∼15 million years ago (Figure 1) thereby creating a large neo-sex chromosome (Carvalho and Clark 2005). The location of the centromere (metacentric or telocentric) prior to fusion is not known, and the A-to-D fusion has been a matter of some debate (Schaeffer 2018). If Muller A was metacentric prior to the fusion, that could explain the presence of ancestral Muller A genes on the long arm of the fused A-D chromosome (denoted XR in *D. pseudoobscura*) (Mahajan et al. 2018, Bracewell et al. 2019) (hereafter referred to as Muller A-AD). Third, the putative Muller C-D fusion is only present in some species of the obscura subgroup, suggesting it occurred recently. How the chromosomes fused is unknown (centromere-centromere, centromere-telomere, telomere-telomere) and the relative size and gene content of this new pericentromeric region is unknown. Here, we report on our genome assembly and annotation of *D. bifasciata* and we characterize chromosome structure, the distribution of transposable elements (TE), and explore the putative Muller C-D fusion.

**Figure 1.**
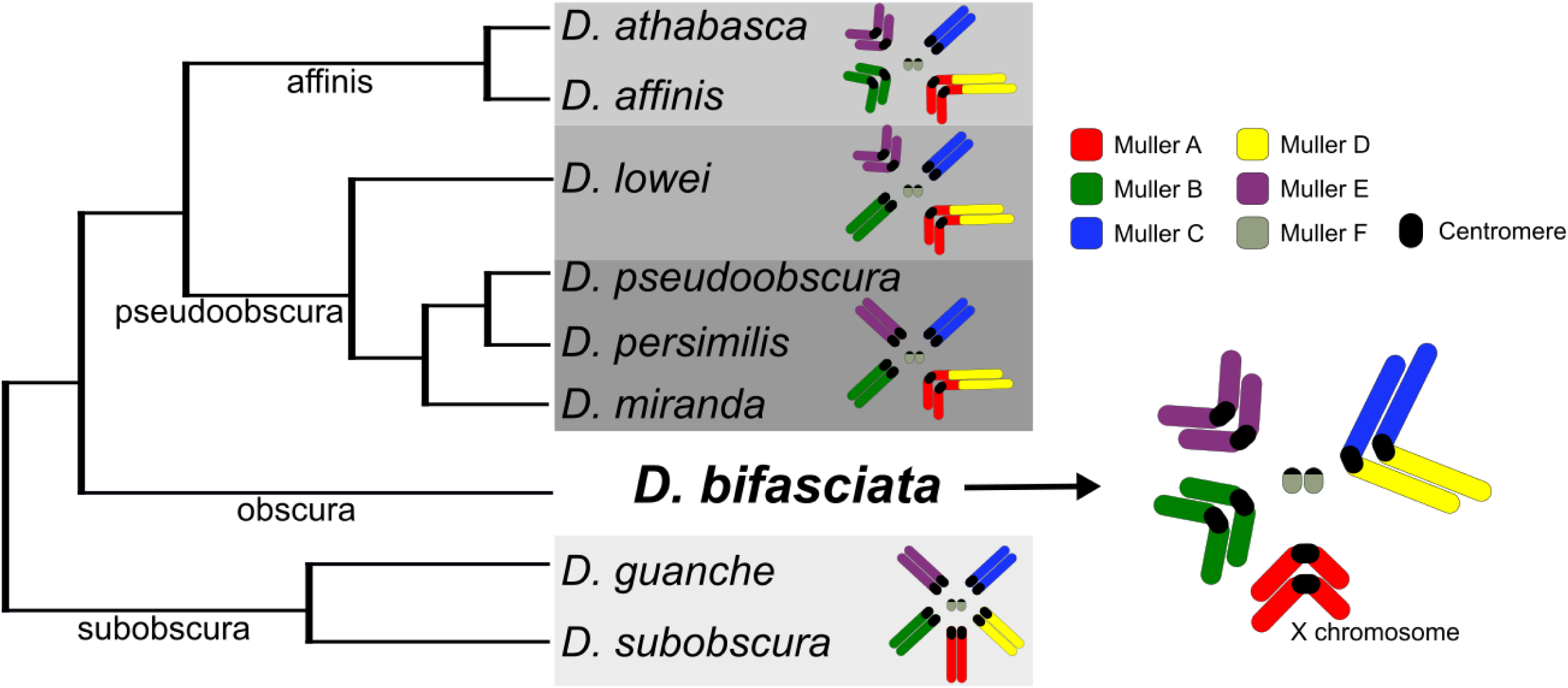
Evolutionary relationships and karyotype transitions of *obscura* group flies. The ancestral karyotype of the *obscura* group (shown here as *Drosophila subobscura*) consists of five large and one small pair of telocentric chromosomes, referred to as Muller elements A-F (reviewed in (Schaeffer 2018)), and shown color coded. Significant karyotypic changes have occurred across the *obscura* group (highlighted with grey boxes) with chromosomal fusions and centromere movement altering chromosome structure (Bracewell et al. 2019). *Drosophila bifasciata* represents an important karyotype to understand evolutionary transitions since Muller A (the X chromosome), B and E are thought to be metacentric and Muller A is unfused (Moriwaki and Kitagawa 1955). In *D. bifasciata*, it is thought that Muller C and D fused, although C-D fusions are only present in some *obscura* subgroup species (Buzzati-Traverso and Scossiroli 1955). Shown phylogenetic relationships adapted from (Gao et al. 2007) with subgroup designations shown along the branches.

## MATERIALS AND METHODS

### Genome sequencing and assembly

We sequenced the *D. bifasciata* isofemale line 14012-0181.02, which was originally collected in Hokkaido, Japan and obtained from the National Drosophila Species Stock Center at Cornell University. High molecular weight DNA for sequencing was extracted from ∼ 60 female flies using a Qiagen Blood & Cell Culture DNA Midi Kit and the resulting DNA was size selected for fragments >15 kb using BluePippin (Sage Science). We generated long-reads using Nanopore and the SQK-LSK109 sequencing kit on one 9.4.1RevD flow cell. Raw output files from our sequencing run were base called using Guppy version 3.0.3 (Oxford Nanopore Technologies) with default parameters for quality score filtering.

We used Canu version 1.8 (Koren et al. 2017) to first error-correct the raw sequencing reads using slightly modified parameters (correctedErrorRate=0.065 corMinCoverage=8 batOptions=“-dg 3 -db 3 -dr 1 -ca 500 -cp 50” trimReadsCoverage=4 trimReadsOverlap=500 genomeSize=200m). The resulting error-corrected reads were then assembled into contigs using the WTDBG2 assembler (Ruan and Li 2019) with default settings. We then BLAST searched all contigs < 1MB to the nt database and returned the top two hits to identify any contigs from non-target species (typically *Acetobacter* and *Saccharomyces*).

After removing contaminant contigs we polished the genome assembly using three rounds of Racon (Vaser et al. 2017) followed by three rounds of Pilon (Walker et al. 2014). This method of combining multiple rounds of Racon and Pilon has been shown to increase genome assembly quality in other Drosophila species (Miller et al. 2018). We used reads derived from our high coverage Hi-C Illumina data (below) for genome polishing. Because of the inherent properties of Hi-C data (paired-end reads with atypical orientations, highly variable insert sizes, chimeric reads) that could lead to spurious genome polishing, we first mapped our Illumina reads to the genome using BWA mem (Li and Durbin 2009) and extracted only those reads with correct pairing using samtools (view -bf 0×2) (Li et al. 2009). We then used those reads as single-end reads for genome polishing. A fraction of the single-end reads will be chimeric but read mapping with BWA mem soft-clips reads and these soft-clipped reads should be randomly distributed across the genome (Figure S1) and not contribute significantly to genome polishing. At each step of assembly and polishing we assessed genome completeness using BUSCO *v3* (Simão et al. 2015) and the odb9 eukaryota database.

### Hi-C scaffolding

Prior to scaffolding we compared our polished contigs with other chromosome-length genome assemblies from *obscura* group species (Mahajan et al. 2018, Bracewell et al. 2019) using whole genome alignments with D-Genies (Cabanettes and Klopp 2018). We then identified the largest contigs belonging to Muller elements to help guide any potential manual manipulations during Hi-C scaffolding. To scaffold the assembly, we used chromatin conformation capture to generate Hi-C data (Lieberman-Aiden et al. 2009). We generated Hi-C libraries as outlined in Bracewell et al. (2019) using a DNase digestion method (Ramani et al. 2016). The resulting DNA library was prepped using an Illumina TruSeq Nano library prep kit and was sequenced on a HiSeq 4000 with 100bp PE reads. We used Juicer (Durand et al. 2016b) to map raw Hi-C reads and generate contact maps based on 3D interactions to scaffold the genome assembly. We then used the 3D-DNA pipeline (Dudchenko et al. 2017) to orient and place contigs. 3D-DNA output files were visualized and checked for accuracy using Juicebox (Durand et al. 2016a) with verification and modifications to scaffolding done using built-in tools. The final assembly was scaffolded together with 300 N’s between each contig.

### Repetitive element identification and genome masking

We first used REPdenovo (Chu et al. 2016) to identify novel repeats from our single-end Hi-C Illumina sequencing data (above) using parameters described in detail in Bracewell et al. (2019). We then concatenated the REPdenovo repeats with the Repbase *Drosophila* repeat library (downloaded March 22, 2016, from www.girinst.org) and used this combined file to mask the genome with RepeatMasker version 4.0.7 using the -no_is and -nolow flags. To characterized the genomic distribution of specific transposable element (TE) families we used a TE library developed from *obscura* group flies (Hill and Betancourt 2018) and again used RepeatMasker and then bedtools coverage (Quinlan and Hall 2010) to determine the proportion of masked bases per TE family.

### Genome annotation and characterization of assembly

To annotate our *D. bifasciata* genome assembly we used the REPdenovo/Repbase repeat-masked genome (above) and the MAKER annotation pipeline (Campbell et al. 2014) to identify gene models. The *ab initio* gene predictors SNAP (Korf 2004) and Augustus (Stanke and Waack 2003) were used to guide the annotation and we used protein sets from *D. pseudoobscura* and *D. melanogaster* (FlyBase) to aid in gene prediction. We used KaryoploteR (Gel and Serra 2017) to plot features of the *D. bifasciata* genome assembly.

### Gene orthologs, genome synteny, and Muller element fusion orientation

To compare our genome assembly with *D. athabasca* which has metacentric Muller A-AD, B and E chromosomes (Figure 1), and *D. subobscura*, which harbors the ancestral karyotype and is composed entirely of telocentric chromosomes (Figure 1), we performed BLASTP reciprocal best hit searches between proteins from our annotations of each species (Bracewell et al. 2019). We used the blast_rbh.py script (Cock et al. 2015) and genomic coordinates of reciprocal best hits were plotted using the *genoPlotR* package (Guy et al. 2010). To determine if the Muller C-D fusion in *D. bifasciata* was the result of a centromere-centromere, centromere-telomere, or telomere-telomere fusion, we identified the 50 most proximal pericentromeric genes from telocentric Muller C and D in *D. subobscura* and plotted the location of orthologs in *D. bifasciata*.

### Data availability

Raw Nanopore and Hi-C (Illumina) reads have been deposited in the NCBI SRA and are under the BioProject (PRJNA565796). The genome assembly and annotation have been deposited with NCBI (accession WIOZ00000000).

## RESULTS AND DISCUSSION

Using one Nanopore flow cell, we generated 538,757 reads that passed Guppy’s standard quality filtering. Our Nanopore reads had an N50 read length of 23,957 bases and provided ∼45× coverage over the genome given an estimated genome size of ∼200Mb for *D. bifasciata*. Our initial hybrid Canu/WTDBG2 assembly resulted in a genome assembly that consisted of 796 contigs with an N50 of 2,325,530. BLAST results flagged multiple putative bacterial contigs (primarily *Acetobacter*) and 49 contigs (5.5Mb of total sequence) were removed. As expected, rounds of Racon polishing (3×) and subsequent Pilon polishing (3×) led to an appreciable increase in our BUSCO scores (Table 1) although the most significant increases in genome completeness were detected after the initial round of Racon or Pilon. Pilon polishing did not lead to as dramatic an increase in genome completeness as seen in other studies (Bracewell et al. 2019) and this was likely due to limitations of our Illumina polishing data that was single-end and was of modest coverage (mean 18×) over the genome (Figure S2). However, we did see a significant increase in genome completeness suggesting that polishing the genome with Hi-C reads can be a viable strategy for increasing genome assembly quality. Our polished genome assembly consisted of 747 contigs with an N50 of 2,386,451. The longest contig was 18,852,285bp with a total genome assembly length of 192,589,718bp.

**Table 1.**
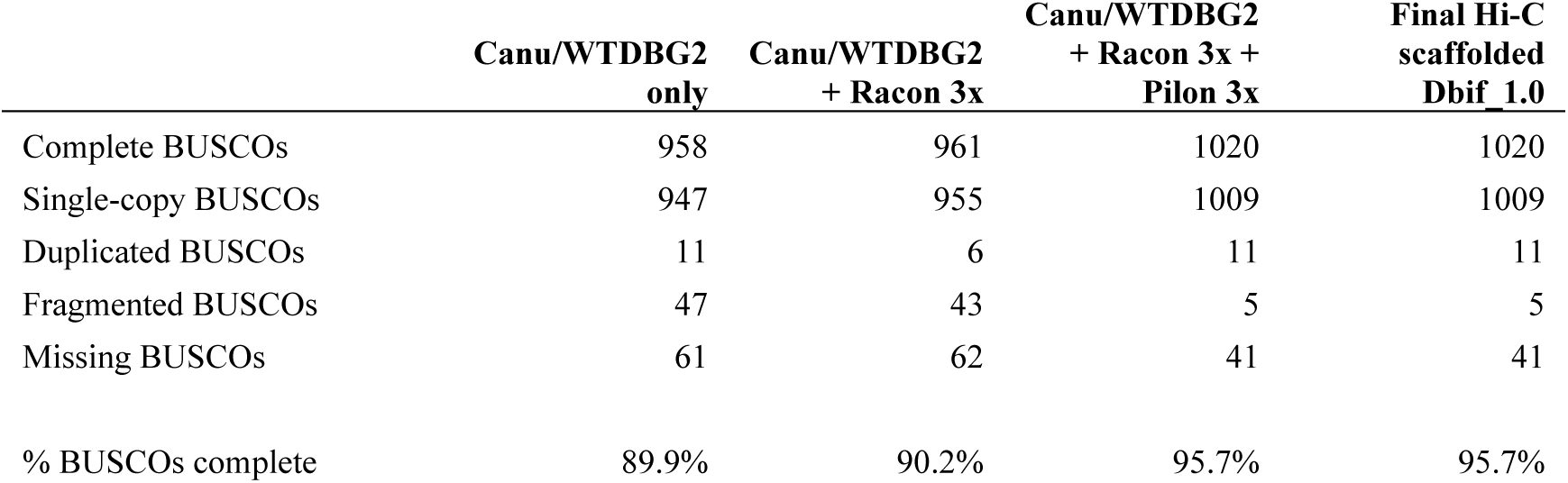
BUSCO results from the genome assembly and polishing process.

Our Hi-C library generated a total of 13,018,415 sequenced read pairs of which 73.8% were alignable to the draft genome. The Juicer pipeline identified 6,734,204 Hi-C contacts which were used to scaffold the genome. The final scaffolded genome assembly placed 126 contigs to the fused Muller CD, 54 to Muller A, 238 to Muller B, 119 to Muller E, 1 to Muller F, and 209 were left unplaced (Figure 2A). Hi-C scaffolding revealed clear associations between euchromatic arms of the same chromosome thereby increasing our confidence in the assembly of metacentric chromosomes (Figure 2A). For example, Muller CD is thought to be the result of a fusion of telocentric Muller C and D elements (Moriwaki and Kitagawa 1955) and our assembly showed clear associations between the C and D arms (Figure 2A). Importantly, there were also clear associations between Muller C and D euchromatic arms with adjacent pericentromeric contigs (Figure 2A), thus providing evidence for the placement of the repeat-rich pericentromeric sequence as well (Figure 2B). Muller A also showed clear associations that extend into highly repetitive pericentromeric regions highlighting this chromosome is indeed metacentric (Figure 2A). The combination of inter-arm and arm-pericentromere Hi-C associations allowed us to determine the correct orientation for all arms of the *D. bifasciata* chromosomes.

BUSCO results suggest our final scaffolded genome assembly is of high quality and 95.7% of BUSCOs were found complete (Table 1). We found the BUSCO statistics to be slightly lower than our other high-quality *obscura* group assemblies which average 98.7% complete (Bracewell et al. 2019). To investigate this reduction, we looked for missing BUSCOs in a species with a similar karyotype and higher score (*D. athabasca*) and found that 49% of missing BUSCOs (20 of 41 total) were in pericentromeric regions. Therefore, residual genome assembly and polishing issues of highly repetitive pericentromeric regions are likely the main contributor to the slightly lower scores of *D. bifasciata*.

**Figure 2.**
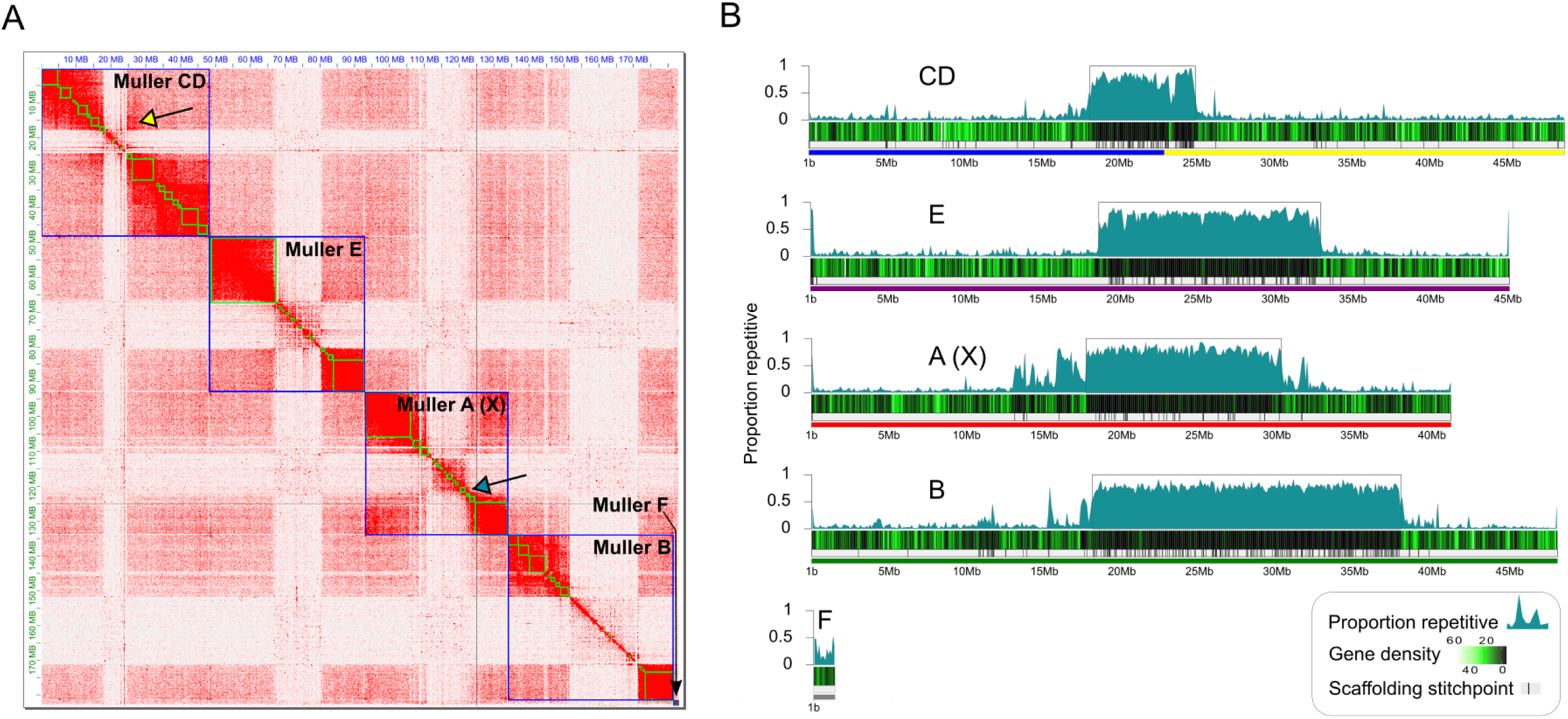
Chromosome-level genome assembly of *Drosophila bifasciata* using Hi-C. A) Hi-C heatmap showing long-range contacts and scaffolding of the genome assembly. Green and blue squares denote contigs and chromosomes, respectively. Euchromatic chromosome arms and heterochromatic pericentromeres for each chromosome show distinct and primarily isolated associations that resemble a ‘checkerboard’ pattern. Note that chromosome arms on opposite sides of a pericentromere often show associations on the diagonal confirming their placement (yellow arrow) while pericentromeres show finer-scale associations with their chromosome arms (blue arrow). B) Shown is the *D. bifasciata* genome assembled into Muller elements (color coded as in Figure 1), scaffolding stitch points, gene density (genes per 100kb) and repeat content (proportion of bases repeat-masked in 100kb non-overlapping windows). Boxes around highly repetitive regions indicate putative pericentromere boundaries (defined as ≥ 40% repeat-masked sequence in sliding windows away from the center).

A total of 57,947,182 bp of the genome assembly was identified as being repetitive (30.1% of the total length of the assembly) and large fractions of all Muller elements were repeat-masked (Figure 2B). The exceptionally high level of repeat-masking located in the middle of chromosome-length scaffolds is indicative of pericentromeric regions on metacentric chromosomes that harbor large numbers of TEs (Kaminker et al. 2002, Bracewell et al. 2019). Indeed, we find that TEs from a few specific families are highly abundant in the pericentromeric region of all Muller elements (Figure 3). Gypsy and Jockey elements are frequently encountered in the pericentromeres of *D. bifasciata* (Figure 3). Nearly 10Mb, and over 5Mb, of assembled sequence (28.4% and 15.8% of all bases masked for TEs) was classified as either Gypsy or Jockey elements, respectively. One specific element, *Daff_Jockey_18*, is at high frequency in all pericentromeres of *D. bifasciata* (Figure S3) and was also the most frequently encountered TE in *D. athabasca*, which also has large metacentric chromosomes (Bracewell et al. 2019).

**Figure 3.**
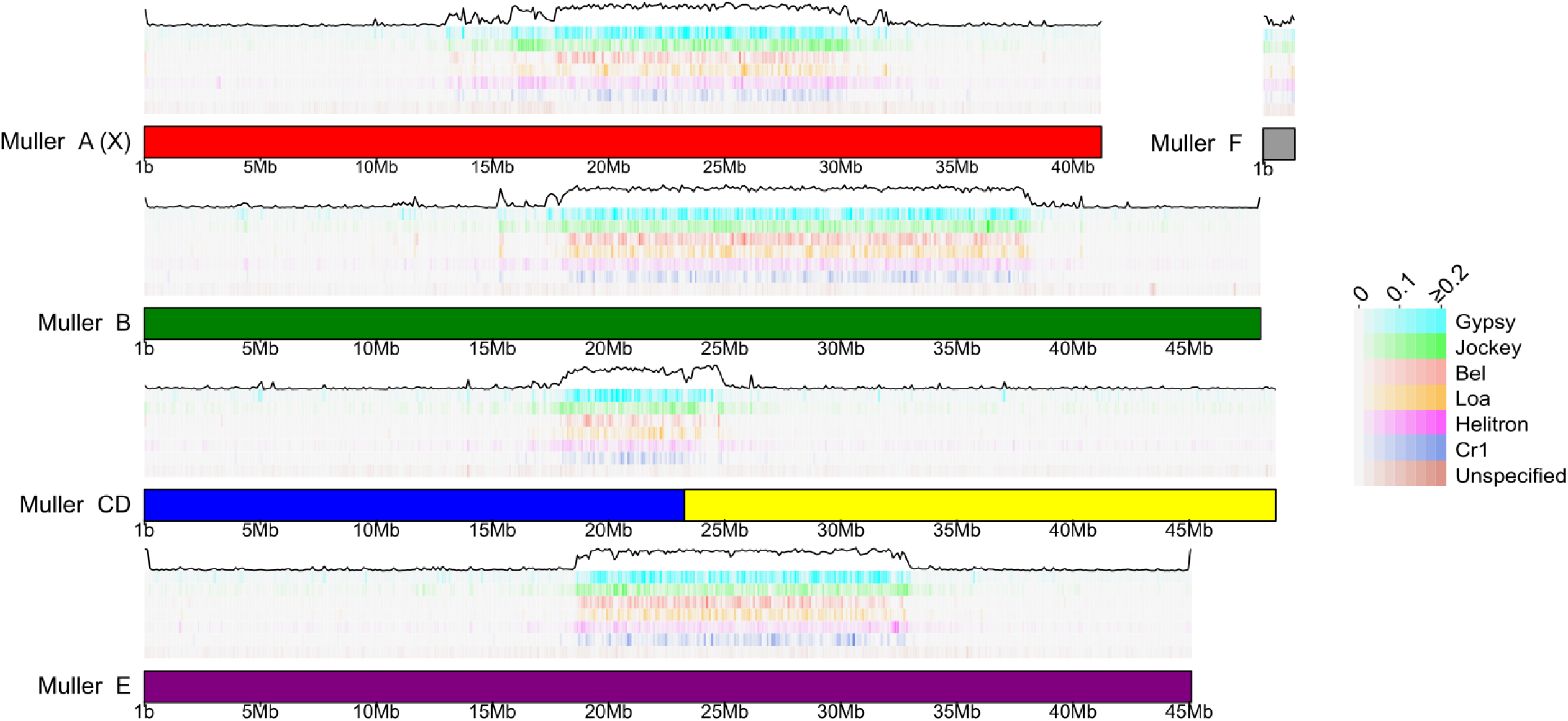
Transposable elements enriched in pericentromeres. Genomic distribution of common transposable element (TE) families in the *D. bifasciata* genome assembly. For each Muller element, TEs are arranged in horizontal tracks of decreasing abundance from top to bottom with the total TE abundance (black line) plotted on top. Shown is the proportion of bases repeat-masked per TE family in 100kb non-overlapping windows.

Our MAKER annotation identified a total of 12,821 protein coding genes models in our *D. bifasciata* genome assembly. This number if very similar to other *obscura* groups species, which have been found to harbor anywhere from 12,714 - 14,547 genes (Mahajan et al. 2018, Puerma et al. 2018, Bracewell et al. 2019, Karageorgiou et al. 2019). We find a total of 2,279 protein-coding genes on Muller A, 2,499 on Muller B, 4,599 on the fused Muller CD, 3,276 on Muller E and 90 on Muller F (Table 2). Comparisons of orthologs between *D. bifasciata* and *D. athabasca* (*affinis* subgroup), which also has a metacentric Muller A-AD, Muller B, and Muller E indicates that the large pericentromeric region in these species is homologous (Figure 4). Surprisingly, the pericentromeric regions in *D. bifasciata* are remarkably similar in size to those of *Drosophila athabasca* suggesting some level of pericentromere stability over long periods of evolutionary time. Conservation of the pericentromere for Muller A between *D. bifasciata* and *D. athabasca* strongly suggests the fusion between Muller A and D involved a telomere-centromere or telomere-telomere fusion between the metacentric Muller A and the telocentric Muller D. This type of fusion would have resulted in the large neo-X (Muller A-AD) we see in species from the *pseudoobscura*/*affinis* subgroup and would account for the excess of Muller A genes on XR of the fused chromosome. For Muller B and Muller E, we find clear evidence of multiple paracentric inversions that differentiate the *D. bifasciata* and *D. athabasca* chromosomes (Figure 4). However, we find no signatures of pericentric inversions, and each arm of Muller B and E appears to be conserved (Figure 4). This pattern contrasts with Muller A where we find evidence of both paracentric and pericentric inversions that differentiate these species (Figure 4).

**Table 2.** Genome assembly and annotation results.

|  | Contigs | Length (bp)<sup>a</sup> | Repetitive<br>(%) | Pericentromere<br>(Mb) | Gene models |
| --- | --- | --- | --- | --- | --- |
| Muller A (X) | 54 | 41,219,968 | 32.4 | 12.6 | 2,279 |
| Muller B | 238 | 48,071,810 | 36.6 | 19.9 | 2,499 |
| Muller CD | 126 | 48,727,904 | 15.8 | 6.8 | 4,599 |
| Muller E | 119 | 45,099,364 | 28.2 | 14.3 | 3,276 |
| Muller F | 1 | 1,364,133 | 26.9 | NA | 90 |
| unplaced | 209 | 8,267,939 | 76.9 | NA | 78 |

<sup>a</sup> Includes N's introduced from scaffolding Muller elements

**Figure 4.**
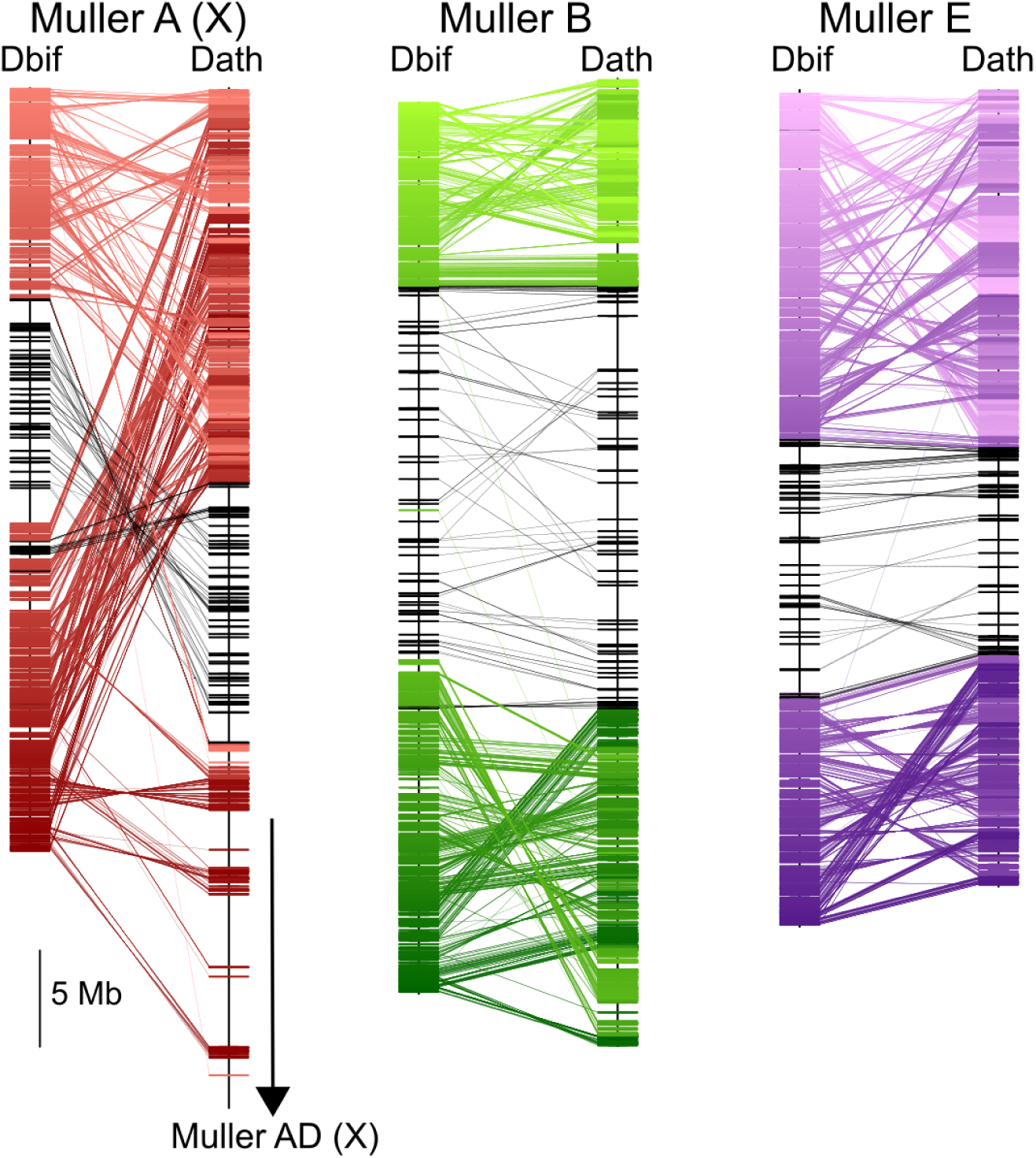
Muller element evolution and synteny. Comparisons of synteny between *D. bifasciata* and *D. athabasca* Muller elements A (X) (red), B (green), and E (purple) with each line representing a protein-coding gene. Genes previously identified as pericentromeric in *D. athabasca* (Bracewell et al. 2019) are shown in black. Only Muller A (X) genes shown for *D. athabasca*.

We also sought to determine the orientation of the fusion between Muller C and D in *D. bifasciata* and we find that the current configuration most likely occurred via a fusion of the two chromosomes at their centromeres. Orthologs of pericentromeric C and D genes in *D. subobscura* are adjacent to one another in our scaffolded assembly (Figure S4) and Hi-C results strongly support this relationship (Figure 2A). Interestingly, the pericentromeric region of the fused C-D chromosome appears smaller than all other pericentromeres in our assembly (Figure 2A). Although speculative, this may be due to the young age of this pericentromere which may be just beginning to expand through the proliferation of repetitive sequences. For example, the 50 pericentromeric C genes in *D. subobscura* are in a 1.0Mb region while orthologs in *D. bifasciata* are spread out across 4.6Mb (Figure S4).

## CONCLUSIONS

Our chromosome-level assembly of *D. bifasciata* provides a valuable resource for future work in this species and will allow for more comprehensive comparative genomic analyses of Drosophila. Our genome assembly method highlights how long-read Nanopore sequencing combined with Hi-C scaffolding can assemble long stretches of highly repetitive pericentromeric sequence, resulting in the assembly of entire metacentric chromosomes. These chromosome-level assemblies allow for evolutionary comparisons of pericentromeric regions that until recently have not been possible. As more chromosome-level genome assemblies become available, we will begin to better understand large-scale changes in chromosome morphology and their impact on genome architecture, gene evolution and speciation.

## ACKNOWLEDGEMENTS

We would like to thank K Wei and D Mai for discussions of repetitive DNA and genome assembly. This work was supported by NIH grants R01GM076007, R01GM101255 and R01GM093182 to D. Bachtrog and 5F32GM123764-02 to R. Bracewell.

## FIGURE CAPTIONS

**Figure S1.**
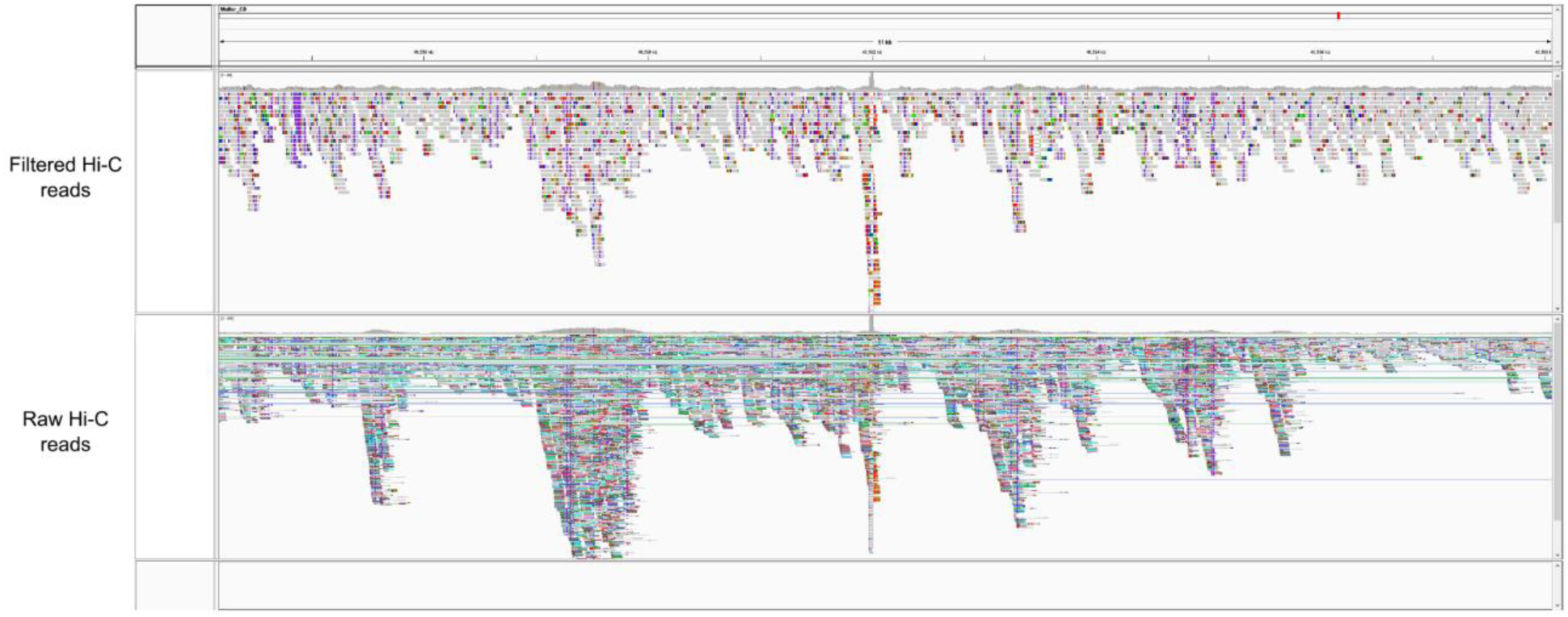
Integrative Genomics Viewer image of a random region of the *D. bifasciata* genome assembly showing mapped raw Hi-C reads (bottom track) and Hi-C reads filtered for Pilon polishing (top track). Raw Hi-C reads are shown as paired (lines connect paired-end reads) and read colors highlight read orientations. Filtered Hi-C reads are shown as single-end with ‘show soft-clipped bases’ enabled. Chimeric reads show up via soft-clipping and can easily be identified since most of the read is aligned (grey) with a portion soft-clipped and shown in alternating red/green/blue. Note the random distribution of soft-clipped reads.

**Figure S2.**
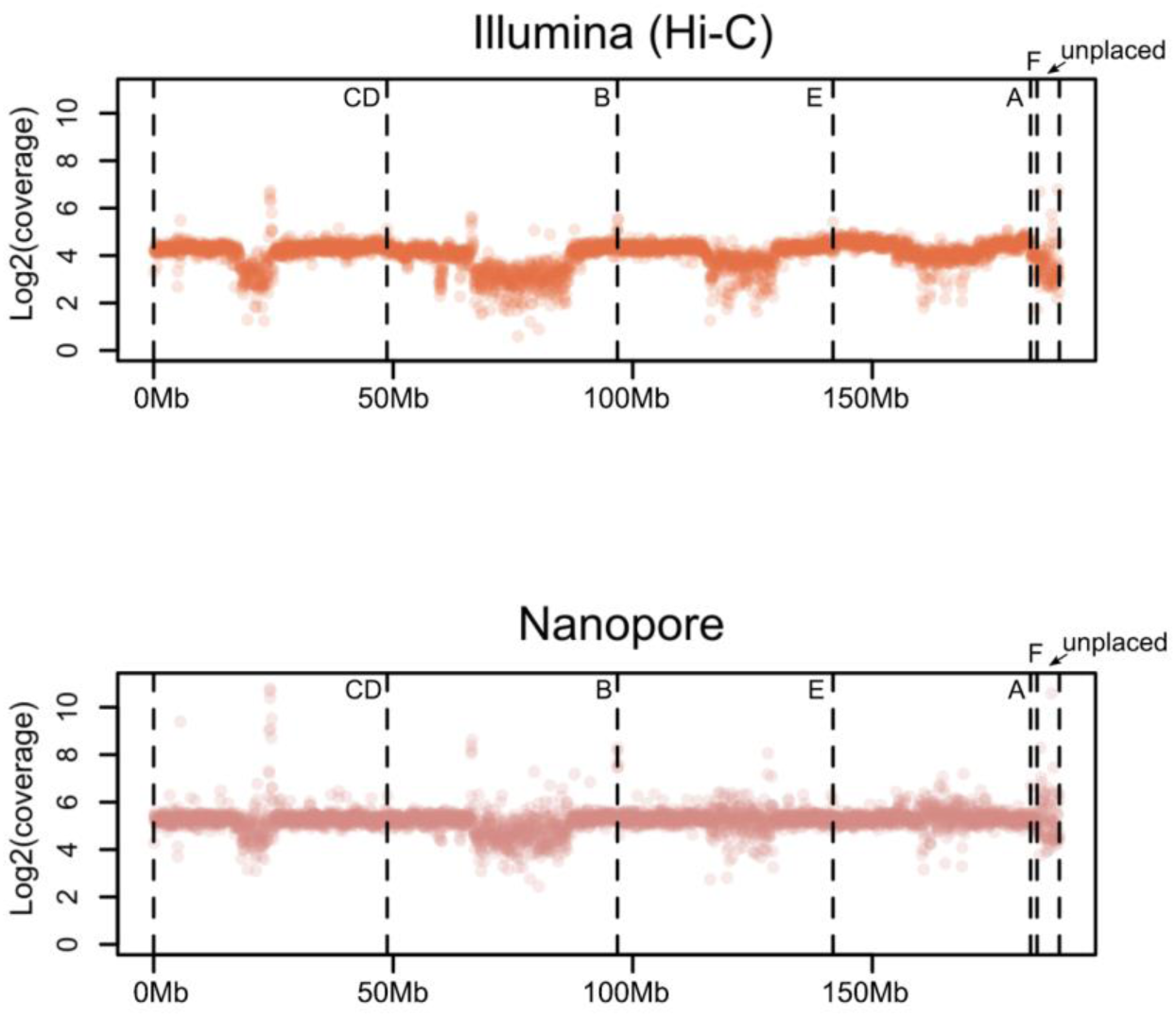
Illumina and Nanopore sequencing coverage over the draft *D. bifasciata* genome assembly.

**Figure S3.**
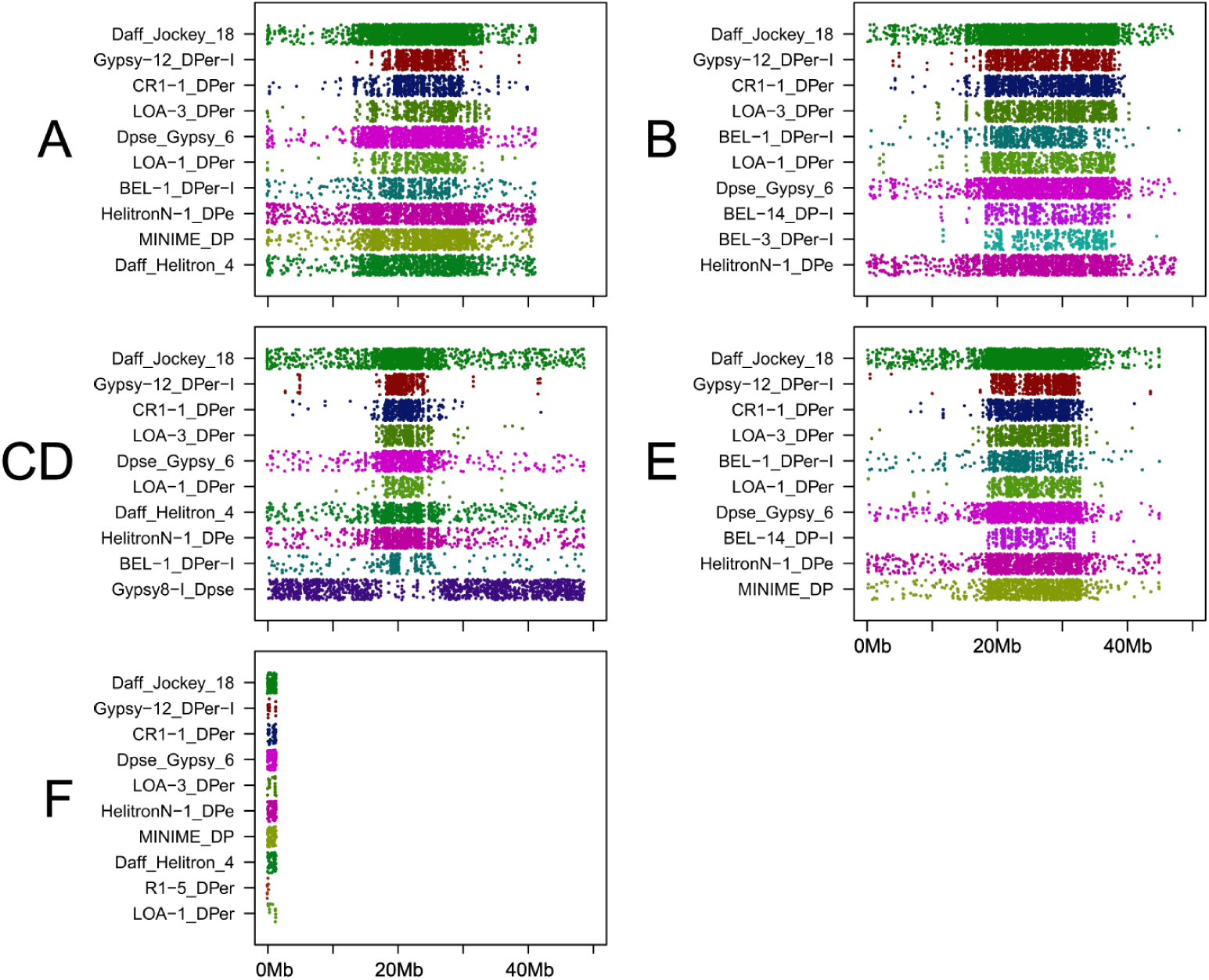
Genomic distribution of the ten most frequently encountered transposable elements (TEs) for each Muller element in the *D. bifasciata* genome assembly. Each point is a genomic location masked for a TE, shown ranked top to bottom by the total amount of masked sequence (bp).

**Figure S4.**
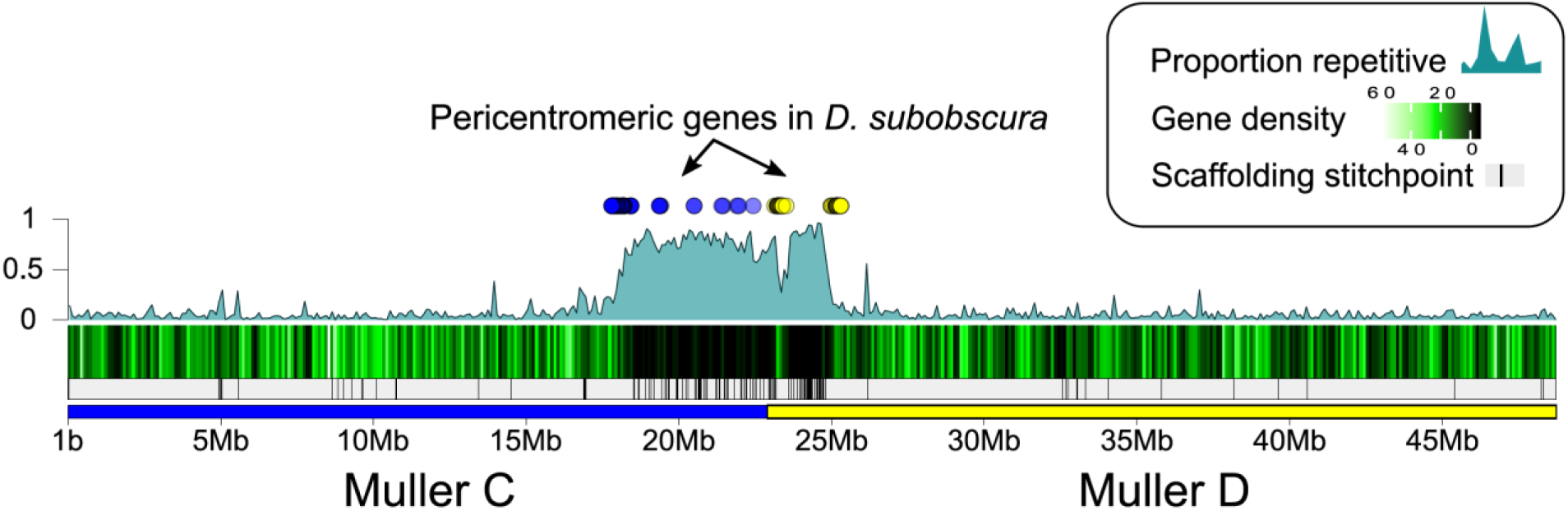
Genomic location of *D. subobscura* pericentromeric orthologs from Muller C (blue dots) and D (yellow dots) on the fused Muller CD of *D. bifasciata*.

